# CreaSol® SSAT (Stabilized Tyrosol) Enhances Creatine on Muscle Performance and Anti-fatigue Capacity in Trained Mice

**DOI:** 10.64898/2026.07.14.738377

**Authors:** Mina Wang, Lily He

## Abstract

Tyrosol, a natural polyphenolic compound, has been identified as a potent cellular antioxidant. It mitigates oxidative stress-induced damage in skeletal muscle cells and facilitates the recovery of intracellular adenosine triphosphate (ATP), suggesting a beneficial role in maintaining cellular energy homeostasis. Creatine, widely used to enhance muscle strength by augmenting the phosphagen system, promotes intracellular ATP production but yields relatively modest improvements in endurance. In this study, we investigated the effects of combined supplementation of tyrosol (CreaSol) and creatine monohydrate (CM) on muscle endurance and strength in mice. Following a 4-week exercise training and intragastric intervention, muscle strength and exercise endurance were evaluated through four consecutive forelimb grip strength tests and exhaustive weighted swimming tests at 24-hour intervals. Compared with creatine monohydrate supplementation alone, the co-administration of CreaSol and creatine monohydrate significantly enhanced grip strength (+28.1%) and swimming endurance (+51.5%). More importantly, following consecutive exhaustive exercise, the combined group exhibited superior recovery capacity, demonstrating significantly attenuated declines in both strength and endurance compared to the creatine monohydrate-only group. We conclude that CreaSol not only effectively improves exercise performance in mice, but also significantly enhances the efficacy of creatine monohydrate in improving muscle strength and endurance, as well as reducing fatigue during consecutive high-intensity exercise.

## Introduction

Enhancing exercise performance and accelerating post-exercise recovery are central topics in sports science and nutrition. As one of the most extensively studied sports supplements, creatine has been well-documented for its efficacy in improving high-intensity exercise capacity, increasing muscle strength, and promoting recovery (1-4). Creatine primarily enhances performance during short-term, high-intensity, and repetitive exercises by facilitating rapid ATP resynthesis. However, numerous studies have demonstrated that creatine supplementation has a less pronounced effect on muscle endurance compared to its impact on muscle strength (5-17). Additionally, current creatine supplementation protocols typically initiate with a loading phase, followed by a daily low-dose maintenance phase to sustain muscle creatine saturation (18-20). Nevertheless, high-dose creatine intake may induce gastrointestinal distress. Conversely, low-dose supplementation alone may be insufficient to elicit ergogenic benefits and has even been deemed ineffective in some studies (21-32). Therefore, identifying natural compounds that can exert synergistic effects with creatine, further enhancing its efficacy or improving the recovery process, holds significant research value and application prospects.

In recent years, certain plant extracts have garnered attention for their antioxidant and anti-fatigue properties. Tyrosol, a phenolic compound found in foods such as olive oil, has been reported to exhibit various biological activities (33-36). It has been confirmed as a potent cellular antioxidant, potentially due to its intracellular accumulation (37). As an effective antioxidant, tyrosol maintains its antioxidant activity even under stringent conditions. Studies have shown that tyrosol can effectively inhibit oxidative damage by partially preventing H_2_O_2_-induced cell death in L6 myotubes through the modulation of extracellular signal-regulated kinase, c-Jun N-terminal kinase, and p38 MAPK pathways, as well as increasing ATP production (38). Furthermore, tyrosol pretreatment has been shown to effectively prevent dexamethasone-induced myotube atrophy and cell death. Its protective mechanism involves maintaining mitochondrial membrane potential and normal morphology, restoring impaired autophagic flux and lysosomal function, and alleviating endoplasmic reticulum stress (39). Given that tyrosol reverses the reduction in ATP production in skeletal muscle under oxidative stress and inhibits hormone-induced skeletal muscle atrophy by maintaining normal mitochondrial function, we hypothesize that tyrosol may promote energy recovery and exert anti-fatigue effects. However, whether tyrosol can enhance creatine to optimize exercise performance and recovery remains unclear.

Based on this, the present study investigated the effects of combined supplementation of tyrosol (CreaSol) and creatine monohydrate on muscle strength, endurance performance, and muscle recovery in mice subjected to consecutive high-intensity exercise. This study aims to provide novel experimental evidence and theoretical support for the development of more efficient, compound sports nutritional supplements.

## Materials and Methods

### Animals

Male Kunming (KM) mice (6 weeks old, SPF grade) were purchased from Shanghai Jiesijie Experimental Animal Co., Ltd. The mice were randomly assigned to groups and housed with four mice per cage. They were acclimatized for 1 week in an SPF-grade animal facility with controlled temperature (20 ± 2°C) and humidity (55 ± 5%), under a 12 h light/dark cycle (lights on from 08:00 to 20:00). Throughout the acclimatization period, the mice had ad libitum access to standard rodent chow and sterilized pure water. Following the final experimental procedure, all mice were anesthetized via intraperitoneal injection of 2.5% Avertin (0.01 mL/g) and subsequently euthanized by cervical dislocation. All animal experimental procedures in this study were strictly conducted in accordance with international guidelines for the welfare of experimental animals, and were reviewed and approved by the Animal Ethics Committee of East China University of Science and Technology.

### Materials

CreaSol^®^ SSAT (Stabilized Tyrosol) and creatine monohydrate were obtained from MolTek Nutrition. The samples were stored in sealed containers, protected from moisture and direct sunlight, with a shelf life of 24 months at room temperature.

### Experimental process

Six-week-old male KM mice were randomly divided into five groups (n = 8): Blank, Control, CreaSol, CM, and CreaSol + CM. The Blank group served as the sedentary blank control, while the Control group acted as the exercise training control. The CreaSol, CM, and CreaSol + CM groups represented the exercise training groups supplemented with the respective test substances.

The experimental protocol lasted for a total of 38 days. Following a 1-week acclimatization period, mice in the CreaSol, CM, and CreaSol + CM groups were intragastrically administered with CreaSol (75 mg/kg/d), creatine monohydrate (260 mg/kg/d), and a combination of both (75 mg/kg/d CreaSol + 260 mg/kg/d creatine monohydrate), respectively. All test substances were dissolved in deionized water, with an administration volume of 10 mL/kg. The mice were weighed weekly, and the daily administration volume was dynamically adjusted to 0.1 mL per 10 g of body weight to ensure accurate dosing. The Blank and Control groups received an equal volume of deionized water. This intervention lasted for 4 weeks. During this period, mice in the Control, CreaSol, CM, and CreaSol + CM groups underwent exercise training 5 min after gavage, consisting of 20 min of unloaded swimming daily, 5 days a week.

On day 35, 30 min after the initial gavage, the first grip strength test was conducted. This was followed by the first exhaustive swimming test after a 1-hour rest. Subsequently, the corresponding samples were administered 24, 48, and 72 h after the first gavage, followed by the second, third, and fourth grip strength and exhaustive swimming tests (Figure 1).

**Figure 1.**
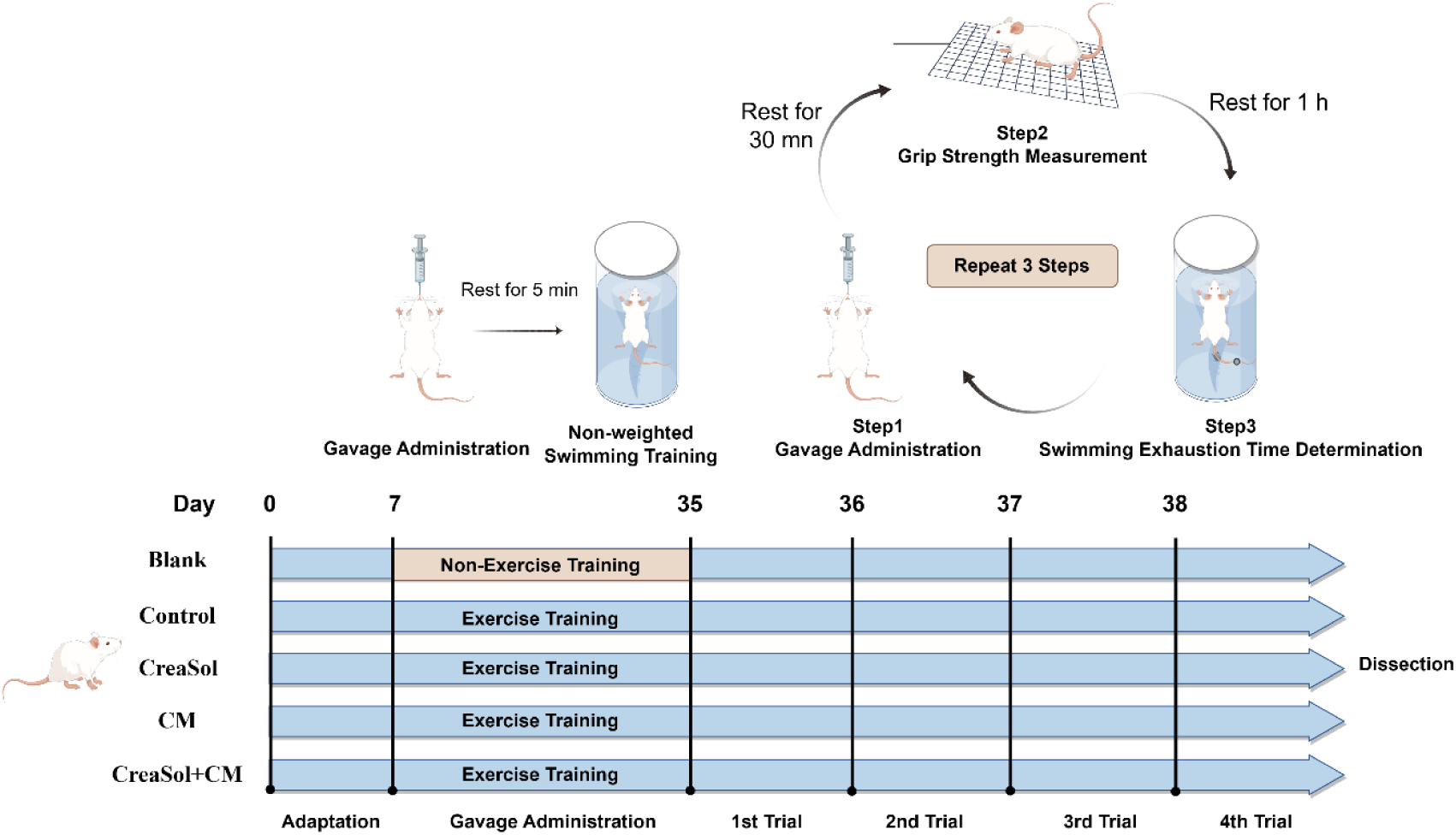
Flow Diagram of Experiment.

### Grip strength test

Forelimb grip strength was measured using a grip strength meter (40, 41). Prior to testing, mice were individually placed in the testing room for a 10–15 min acclimatization period to minimize the impact of stress on muscle performance. The experimenter grasped the base of the mouse’s tail in a stable and consistent manner, keeping the body horizontal, and gently guided the mouse toward the metal grid at the front of the apparatus to allow a natural grip. Once the mouse firmly grasped the grid with all four limbs, the experimenter pulled the mouse backward along its horizontal axis at a constant speed until the grip was broken. The tension sensor within the grip strength meter recorded the peak pull force exerted by the mouse before releasing the grid, and the results were displayed in gram-force (gf). Each mouse was tested consecutively three times, with approximately a 1-min interval between trials to prevent fatigue. The average of the three trials was calculated and used as the final grip strength value for each animal. To ensure data reliability, all tests were conducted by the same experimenter during a fixed time of day, and the grip strength meter was calibrated before each testing session.

### Exhaustive swimming test

The weighted swimming test was employed to evaluate the endurance and anti-fatigue capacity of the mice (42). Prior to the test, tin wire equivalent to approximately 5% of the mouse’s body weight was evenly wrapped and secured around the proximal part of the tail to provide a stable additional load during swimming. The mouse was then gently placed into a pre-prepared swimming tank. The water temperature was maintained at 30 ± 1 °C, and the water depth was set to prevent the mouse from touching the bottom and standing, thereby ensuring continuous swimming. Throughout the procedure, the experimenter continuously monitored the behavioral status of the mouse. Exhaustion was defined as touching the bottom three consecutive times within 30 seconds, indicating a clear inability to maintain a normal swimming posture and a reliance on the pool bottom for support. Upon reaching exhaustion, the mouse was immediately removed, dried, and placed on a heating pad to restore body temperature. The time from entering the water to exhaustion (time to exhaustion) was recorded for each mouse as an indicator of endurance.

## Results

### CreaSol Enhances the Efficacy of Creatine Monohydrate on Muscle Strength

Based on the grip strength test results (Figure 2), although exercise training significantly increased the grip strength of mice from 230.6 ± 18.3 gf to 339.3 ± 36.0 gf, creatine monohydrate and the combined supplementation exhibited a more pronounced enhancement, with grip strengths reaching 408.1 ± 25.9 gf and 407.6 ± 10.8 gf, respectively (Figure 2A).

**Figure 2.**
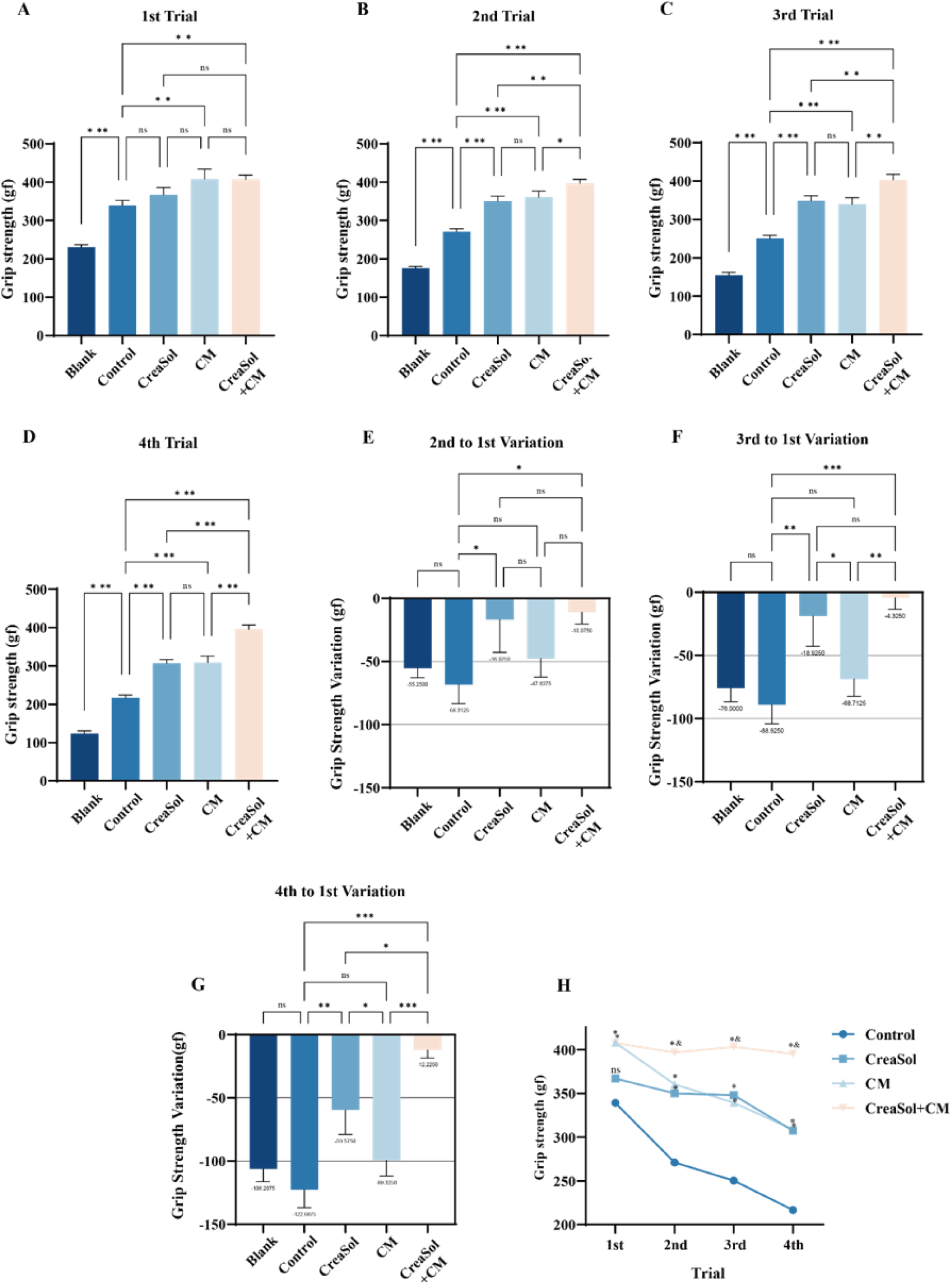
Effects of CreaSol combined with creatine monohydrate on the grip strength in mice. A-D: Grip strength of mice in the 1st to 4th tests. E-G: Grip strength in each test expressed as a percentage relative to the 1st test. H: Changes in grip strength across the tests in the four groups. In panels A-G, * indicates significant differences between groups (*p < 0.05, **p < 0.01, ***p < 0.001); ns indicates no significant difference. In panel H, * indicates a significant difference compared to the Control group (p < 0.05), and & indicates a significant difference compared to the CM group (p < 0.05); ns indicates no significant difference (p > 0.05). Data are presented as Mean ± SEM.

During the three recovery assessments conducted at 24-hour intervals post-exhaustion (Figure 2B-G), the CreaSol group and combined group (CreaSol + CM) showed a significantly lower degree of grip strength decline compared to the control group. A comprehensive analysis of the grip strength variation curves across the four tests (Figure 2H) revealed that, despite similar initial grip strengths between the CM and combined groups, their performance following consecutive exhaustion differed significantly. The grip strength of the CM group dropped markedly after the first exhaustion, and its recovery capacity significantly deteriorated as the number of exhaustive bouts increased; the grip strengths in the subsequent three tests recovered to only 89.24%, 83.81%, and 76.1% of the initial value, respectively. In contrast, the combined group exhibited superior anti-fatigue properties with almost no decline in grip strength, recovering to 97.52%, 98.94%, and 97.04% of the initial value in the subsequent three tests, respectively. The recovery effect of the CreaSol group was slightly inferior to that of the combined group, with recovery rates of 97.4%, 96.55%, and 85.05% in the subsequent three tests. In the fourth test, compared to creatine monohydrate supplementation alone, the combined supplementation increased muscle strength by 28.1%, and by 82.5% compared to the control group.

### CreaSol Enhances the Efficacy of Creatine Monohydrate on Muscle Endurance and Anti-Fatigue in Trained Mice

The results of the swimming exhaustion test (Figure 3) showed that although exercise training alone extended the time to exhaustion of mice from 326.0 ± 66.07 s to 421.5 ± 53.93 s, the difference did not reach statistical significance (p > 0.05). Similarly, supplementation with creatine monohydrate alone showed a certain improving trend (590.0 ± 90.64 s), but it also lacked statistical significance (p > 0.05). In contrast, the CreaSol group and the combined supplementation group exhibited a significant promoting effect on extending the time to exhaustion, reaching 672.5 ± 69.81 s and 768.9 ± 83.1 s, respectively (Figure 3A).

**Figure 3.**
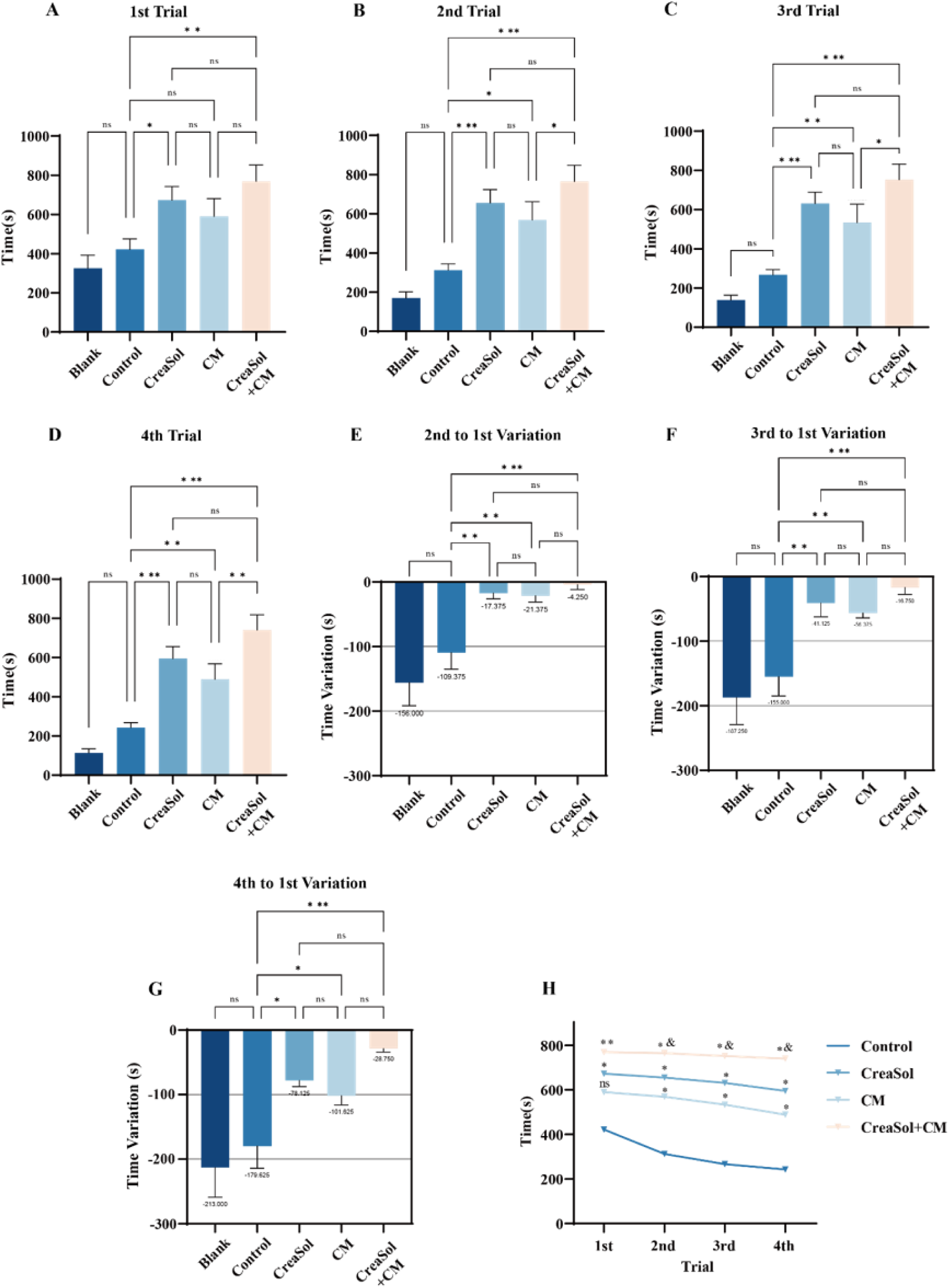
Effects of CreaSol combined with creatine monohydrate on the time to exhaustion in mice. A-D: Time to exhaustion of mice in the 1st to 4th tests. E-G: Time to exhaustion in each test expressed as a percentage relative to the 1st test. H: Changes in time to exhaustion across the tests in the four groups. In panels A-G, * indicates significant differences between groups (*p < 0.05, **p < 0.01, ***p < 0.001); ns indicates no significant difference. In panel H, * indicates a significant difference compared to the Control group (p < 0.05), and & indicates a significant difference compared to the CM group (p < 0.05); ns indicates no significant difference (p > 0.05). Data are presented as Mean ± SEM.

Further analysis of the recovery status after three consecutive exhaustive swimming bouts at 24-hour intervals (Figure 3B-G) revealed that the recovery trend of swimming time to exhaustion across groups was highly consistent with the grip strength test results. Compared to the exercise-only group, the recovery capacity of mice in all three intervention groups was improved. In particular, during the subsequent consecutive exhaustive tests, the combined group demonstrated superior anti-fatigue and endurance-maintaining capabilities, effectively mitigating the decline in exercise performance. Compared to the initial time to exhaustion, the combined group exhibited the smallest decline across the four tests (Figure 3E-G). In the fourth exhaustive swimming test, the time to exhaustion in the combined group was significantly extended by 205.4% compared to the control group, and by 51.5% compared to the CM group.

From an intra-group comparison, the time to exhaustion in the fourth exhaustive swimming test for the combined group decreased by only 3.54% compared to the first test, demonstrating excellent endurance stability. This was followed by the CreaSol group, which showed a 11.55% decrease, while the CM group exhibited a 18.12% decrease. A comprehensive analysis of the variation curves for the time to exhaustion across the four tests (Figure 3H) clearly indicated that the combined group maintained a significantly longer exhaustion duration at each time point compared to the other groups, with an extremely gradual overall declining trend. Statistical analysis further confirmed that the performance of the combined group in the 2nd to 4th exhaustive swimming tests was significantly superior to that of the CM group. In summary, the combination of CreaSol and creatine monohydrate exhibits the optimal synergistic effect in improving muscle endurance and reducing fatigue.

## Discussion

As one of the most extensively studied sports nutritional supplements, the mechanisms by which creatine enhances exercise performance have been widely studied. Creatine primarily augments the efficiency of the phosphagen system by increasing intramuscular phosphocreatine stores and accelerating ATP resynthesis. This metabolic adaptation yields significant ergogenic benefits for short-duration, high-intensity explosive exercises and resistance training. However, the benefits of creatine supplementation on endurance performance remain controversial. Multiple studies have pointed out that creatine alone does not significantly improve aerobic endurance or prolonged continuous exercise. In some cases, creatine-induced weight gain (due to water retention) may even negatively impact endurance events. Furthermore, traditional creatine supplementation protocols typically require a “loading phase,” involving short-term high-dose intake, which is often accompanied by gastrointestinal distress, nausea, and bloating, thereby reducing user compliance. Therefore, exploring a synergistic strategy that can reduce the required dosage of creatine, mitigate its side effects, and broaden its ergogenic benefits— especially in the realm of endurance—holds significant scientific and practical value.

In this study, we introduced the natural plant-derived bioactive compound tyrosol to investigate its potentiating effects on creatine efficacy. The results from our animal experiments reveal the unique advantages of combining tyrosol (CreaSol) with creatine in improving exercise performance, particularly their synergistic effects in maintaining muscle strength and endurance. Regarding muscle strength, the combined supplementation significantly promoted the recovery of grip strength post-exhaustion in mice. Notably, compared to creatine supplementation alone, the combined group demonstrated greater efficacy in maintaining strength output during consecutive exercise, significantly delaying the decline in muscle strength. This finding suggests that CreaSol may optimize creatine utilization or the muscle energy metabolic environment through certain mechanisms, thereby maintaining or even surpassing the strength-enhancing effects of creatine alone.

More importantly, our study also uncovered the synergistic potential of CreaSol in enhancing endurance performance, which to some extent breaks through the limitations of traditional creatine supplementation. In the swimming exhaustion test, the combination of tyrosol and creatine not only significantly prolonged the time to exhaustion in mice but also effectively maintained endurance levels during consecutive exhaustive exercise, demonstrates excellent fatigue resistance. This indicates that the addition of CreaSol not only preserves the benefits of creatine on the phosphagen system but may also compensate for the shortcomings of creatine alone in aerobic endurance through other metabolic pathways, such as antioxidant activity and the improvement of mitochondrial function. This dual “strength-endurance” enhancing property makes this combination highly promising for addressing complex, mixed-modality exercise demands.

The present study demonstrates that the natural plant-derived compound CreaSol significantly enhances the efficacy of creatine. The combination of CreaSol not only maintains the high efficacy of creatine in enhancing muscle strength, but more importantly, it significantly broadens the efficacy of creatine in improving muscle endurance and anti-fatigue capability. This finding provides robust experimental evidence for the development of novel, low-side-effect, and comprehensive sports nutritional supplements, offering new insights to address the issues of poor tolerability and limited endurance benefits associated with traditional creatine supplementation.

## Acknowledgements

We are deeply grateful to Fangci Xiong from East China University of Science and Technology for her valuable technical support. This study was sponsored by MolTek Nutrition Co., Ltd.

